# Direct nucleic acid analysis of mosquitoes for high fidelity species identification and detection of *Wolbachia* using a cellphone

**DOI:** 10.1101/291849

**Authors:** Sanchita Bhadra, Timothy E. Riedel, Miguel A. Saldaña, Shivanand Hegde, Nicole Pederson, Grant L. Hughes, Andrew D. Ellington

## Abstract

Manipulation of natural mosquito populations using the endosymbiotic bacteria *Wolbachia* is being investigated as a novel strategy to reduce the burden of mosquito-borne viruses. To evaluate the efficacy of these interventions, it will be critical to determine *Wolbachia* infection frequencies in *Aedes aegypti* mosquito populations. However, current diagnostic tools are not well-suited to fit this need. Morphological methods cannot identify *Wolbachia*, immunoassays often suffer from low sensitivity and poor throughput, while PCR and spectroscopy require complex instruments and technical expertise, which restrict their use to centralized laboratories. To address this unmet need, we have used loop-mediated isothermal amplification (LAMP) and oligonucleotide strand displacement (OSD) probes to create a one-pot sample-to-answer nucleic acid diagnostic platform for vector and symbiont surveillance. LAMP-OSD assays can directly amplify target nucleic acids from macerated mosquitoes without requiring nucleic acid purification and yield specific single endpoint yes/no fluorescence signals that are observable to eye or by cellphone camera. We demonstrate cellphone-imaged LAMP-OSD tests for two targets, the *Aedes aegypti* cytochrome oxidase I (*coi*) gene and the *Wolbachia* surface protein (*wsp*) gene, and show a limit of detection of 4 and 40 target DNA copies, respectively. In a blinded test of 90 field-caught mosquitoes, the *coi* LAMP-OSD assay demonstrated 98% specificity and 97% sensitivity in identifying *Ae. aegypti* mosquitoes even after 3 weeks of storage without desiccant at 37 °C. Similarly, the *wsp* LAMP-OSD assay readily identified the *w*AlbB *Wolbachia* strain in field-collected *Aedes albopictus* mosquitoes without generating any false positive signals. Modest technology requirements, minimal execution steps, simple binary readout, and robust accuracy make the LAMP-OSD-to-cellphone assay platform well suited for field vector surveillance in austere or resource-limited conditions.

**Author summary:** Mosquitoes spread many human pathogens and novel approaches are required to reduce the burden of mosquito-borne disease. One promising approach is transferring *Wolbachia* into *Aedes aegypti* mosquitoes where it blocks transmission of arboviruses like dengue, Zika and Yellow fever viruses and spreads through mosquito populations. For effective evaluation of this approach, regular surveillance of *Wolbachia* infections in *Ae. aegypti* is required, but current diagnostic tools are not well suited to support these critical surveillance needs. To fill this need we developed a simple, robust and inexpensive assay to identify *Ae. aegypti* mosquitoes and *Wolbachia* using our unique one-pot assay platform, LAMP-OSD, which uses loop-mediated isothermal amplification to amplify nucleic acid targets at a single temperature. Unlike other LAMP-based tests, our assays assure accuracy by coupling amplification with novel nucleic acid strand displacement (OSD) probes that hybridize to specific sequences in LAMP amplification products and thereby generate simple yes/no readout of fluorescence readable by human eye and by off-the-shelf cellphones. To facilitate field use, we developed our assays so they are compatible with crushed mosquito homogenate as the template, meaning no nucleic acid extraction is required. In blinded tests using field collected mosquitoes, LAMP-OSD-cellphone tests performed robustly to identify 29 of 30 *Ae. aegypti* even after 3 weeks of storage at 37 °C while producing only one false positive out of 60 non-specific mosquitoes. Similarly, our assay could identify *Wolbachia* in field-caught *Aedes albopictus* without producing any false positives. Our easy to use and easy to interpret assays should facilitate widespread field mosquito surveillance with minimal instrumentation and high accuracy.

## Introduction

Mosquitoes are vectors that can transmit an array of pathogens that often cause devastating human diseases [1]. Traditionally considered a problem for tropical regions, mosquitoes are increasingly becoming a global public health challenge [2, 3] due to a changing global environment, urbanization, increases in the global movement of populations, and the emergence of insecticide resistance [4]. Estimates suggest nearly half the world’s population is at risk for mosquito-borne diseases [5, 6], and as such, there is an urgent need for novel approaches to reduce the burden of disease.

One biocontrol countermeasure gaining traction for mosquito control is the release of *Wolbachia* infected mosquitoes [7–9]. *Wolbachia* is a maternally-transmitted endosymbiont that can rapidly become established in the natural mosquito populations and can inhibit a variety of pathogens, including arboviruses, malaria parasites, and filarial nematodes [10–15]. *Wolbachia* control strategies are currently being deployed into the field to alter the capacity of *Aedes aegypti* to transmit arboviruses or to suppress mosquito populations [16–18]. Surveillance of transinfected mosquitoes as well as natural vector populations is crucial to evaluate the efficacy of these interventions [19]. However, most current screening methods rely on PCR, which is expensive and relies on laboratory facilities. In addition to screening for *Wolbachia* infection, it would also be desirable to identify the host mosquito species using these assays since different mosquito species differ in their ability to transmit pathogens [20]. Knowledge of vector species, and prevalence and stability of *Wolbachia* is essential for effective vector control and pre-emption of disease outbreaks with public health measures [21].

Unfortunately, mosquitoes are most commonly identified using morphological taxonomic keys. This process can be tedious, and requires highly trained personnel and undamaged mosquitoes. Alternative morphological methods such as the identification of morphometric wing characters [22] are low throughput and require microscopes and complex imaging instruments. Moreover, traditional morphological-based approaches cannot detect associated symbionts or pathogens. These limitations restrict widespread accessibility and necessitate sample preservation and transport. On the other end of the spectrum, immunoassay-based tools for identifying pathogen-infected mosquitoes, such as VecTest™ dipsticks (Medical Analysis Systems Inc.), are portable and inexpensive. However, these tests have poor sensitivity [23–25] and are not necessarily available for distinguishing mosquito species or identifying *Wolbachia* endosymbionts.

Nucleic acid tests can provide the necessary sensitivity and versatility for identifying both *Wolbachia* and mosquito species. However, since molecular testing is currently heavily reliant on PCR [26–28], opportunities for field-based determinations are limited, leading to significant delays and gaps in actionable surveillance. To support widespread vector surveillance inexpensive, portable, nucleic acid diagnostic platforms are needed that rapidly produce accurate results without requiring complex procedures, instruments, and laboratory infrastructure. In this regard, isothermal nucleic acid amplification assays such as loop-mediated isothermal amplification (LAMP) have begun to be employed because they do not require complex thermocycling instruments [29–31]. However, although LAMP can rival PCR for sensitivity it often produces spurious amplicons, which in turn lead to false positive readouts with non-specific reporters such as Mg^2+^ precipitation or fluorescent dye intercalation [32–34].

To mitigate the spurious signals that arise with LAMP, we have previously applied principles that were developed for nucleic acid strand exchange circuits [35–38] to the design of short hemiduplex oligonucleotide strand displacement (OSD) probes for LAMP [39]. The single stranded ‘toehold’ regions of OSD probes bind to LAMP amplicon loop sequences, and then signal via strand exchange [40] that leads to separation of a fluorophore and quencher [39]. OSDs are the functional equivalents of TaqMan probes and can specifically report single or multiplex LAMP amplicons without interference from non-specific nucleic acids or inhibitors [39, 41]. OSDs significantly enhance the diagnostic applicability of LAMP, allowing it to match the allelic specificity of real-time PCR. Recently, we engineered these molecular innovations to function fluently in one-pot LAMP-OSD reactions that can directly amplify a few tens to hundreds of copies of DNA and RNA analytes from minimally processed specimens and produce sequence-specific fluorescence signals that are easily observable by the human eye or (more importantly) by unmodified cellphone cameras [42]. The fluorescence endpoints that are produced can be used for yes/no determinations of the presence of an analyte, and also estimation of analyte copies on an order of magnitude scale [42, 43].

Here, we have adapted our smartphone-read one-pot LAMP-OSD system to directly amplify target nucleic acids from crudely macerated mosquitoes and to sequence-specifically report both mosquito and symbiont amplicons as visually readable fluorescence. In particular, we have developed two LAMP-OSD assays – one targeting the *Ae. aegypti* cytochrome oxidase I gene (*coi*), and the other the *Wolbachia w*AlbB surface protein (*wsp*) gene. Using a blinded set of field-caught mosquitoes, we demonstrate the exquisite sensitivity and specificity of our LAMP-OSD platform for identifying mosquito species and detecting *Wolbachia* infections.

## Methods

### Chemicals and reagents

All chemicals were of analytical grade and were purchased from Sigma-Aldrich (St. Louis, MO, U.S.A.) unless otherwise indicated. All enzymes and related buffers were purchased from New England Biolabs (NEB, Ipswich, MA) unless otherwise indicated. All oligonucleotides and gene blocks (summarized in **S1 Table)** were obtained from Integrated DNA Technologies (IDT, Coralville, IA, U.S.A.).

### Preparation of amplification targets

*Ae. aegypti coi* LAMP-OSD target sequence was amplified by PCR from mosquito genomic DNA. LAMP-OSD target region of the *Wolbachia w*AlbB *wsp* gene was purchased as a gBlock fragment. Both amplification targets were cloned into the pCR2.1-TOPO vector (Fisher Scientific, Hampton, NH) by Gibson assembly according to manufacturer’s (NEB) instructions [44]. Cloned plasmids were selected and maintained in an *E. coli* Top10 strain. Plasmid minipreps were prepared from these strains using the Qiagen miniprep kit (Qiagen, Valencia, CA, USA). All target inserts were verified by sequencing at the Institute of Cellular and Molecular Biology Core DNA Sequencing Facility.

### LAMP primer and OSD probe design

*Wolbachia w*AlbB and *w*Pip strain *wsp* genes and *Ae. aegypti coi* gene sequences were obtained from NCBI GenBank. Consensus signature sequences were derived following MUSCLE (MUltiple Sequence Comparison by Log-Expectation) alignment analysis of each gene set. Target specificity of these signature sequences was evaluated by comparing them to respective *wsp* or *coi* gene sets from phylogenetically-related strains and species such as *Wolbachia w*Mel and *Ae. albopictus*, respectively. Both MUSCLE alignment as well as NCBI BLAST [45, 46] analysis were used for this *in silico* specificity analysis.

The Primer Explorer v5 primer design software (Eiken Chemical Co., Japan) was used for generating several potential LAMP primer sets composed of the outer primers F3 and B3 and the inner primers FIP and BIP. Primer design was constrained to include at least a 40 bp gap between the F1 and F2 or between the B1 and B2 priming sites. Primer specificity for targeted sequence and a corresponding lack of significant cross-reactivity to other nucleic acids of human, vector or pathogenic origin were further assessed using NCBI BLAST.

These primer sets were functionally tested in LAMP assays using zero to several hundred copies of purified plasmids as templates. Amplification kinetics were measured in real time using the fluorogenic intercalating dye Evagreen and the LightCycler 96 real-time PCR machine (Roche). The fastest primer sets that detected the fewest template copies with negligible spurious reactivity in the absence of templates were selected for further assay development.

Fluorogenic OSD probes were then designed to undergo toehold-mediated strand exchange with these *Ae. aegypti coi* and the *Wolbachia wsp* LAMP amplicons. Of the two target derived loop regions (between the F1c and F2, and the B1c and B2 primer binding sites) the regions between F1c and F2 were chosen as *wsp* and *coi* OSD binding regions. Fluorogenic OSD probes were designed using the NUPACK software and our previously described engineering principles [39]. Briefly the hemiduplex OSDs were designed to display 11-12 nucleotide long single-stranded toeholds on the longer, fluorophore-labeled strands. All free 3’-OH ends were blocked with inverted dT to prevent extension by DNA polymerase.

Single loop primers were designed to bind the second loop region (between B1c and B2 primer binding sites) of the *wsp* and *coi* LAMP amplicons and accelerate LAMP amplification.

### LAMP assay with synthetic DNA template

LAMP assays were assembled in a total volume of 25 µl of 1X Isothermal buffer (NEB; 20 mM Tris-HCl, 10 mM (NH_4_)_2_SO_4_, 50 mM KCl, 2 mM MgSO_4_, 0.1% Tween 20, pH 8.8 at 25°C). The buffer was supplemented with 0.4 mM dNTPs, 0.8 M betaine, 2 mM additional MgCl_2_, 1.6 µM each of FIP and BIP, 0.8 µM of loop primer, 0.4 µM each of F3 and B3 primers, and 16 units of *Bst* 2.0 DNA polymerase. Plasmid DNA templates were serially diluted in TE buffer (10 mM Tris-HCl, pH 7.5:0.1 mM EDTA, pH 8.0) immediately prior to use. Zero to several hundred copies of synthetic plasmid templates were added to the LAMP reaction mixes followed by 90 min of incubation at 65 °C.

### Real-time detection of LAMP amplicons using intercalating dyes

1X EvaGreen (Biotium, Hayward, CA, USA) was included in LAMP assays that were then analyzed using the LightCycler 96 real-time PCR machine (Roche, Basel, Switzerland). Reactions were subjected to 45 cycles of two-step incubations – step 1:150 sec at 65 °C, step 2: 30 sec at 65 °C. EvaGreen signal was measured in the FAM channel during step 2 of each cycle. Subsequently, amplicons were subjected to a melt analysis by incubation at 65 °C for 1 min followed by incremental rise in temperature to 97 °C. Amplicon melting was monitored by measuring fluorescence at the rate of 10 readings per °C change in temperature. The resulting data was analyzed using the LightCycler 96 analysis software to measure Cq values for amplification and amplicon melting temperatures.

### Real-time detection of LAMP amplicons using OSD probes

LAMP reactions monitored in real time using OSD probes were assembled and analyzed as above with the following changes. First, OSD probes were prepared by annealing 1 µM of the fluorophore-labeled OSD strand with 5 µM of the quencher-labeled strand in 1X Isothermal buffer. Annealing was performed by denaturing the oligonucleotide mix at 95 °C for 1 min followed by slow cooling at the rate of 0.1 °C/s to 25 °C. Excess annealed probe was stored at −20 °C. Annealed OSD probes were added to the LAMP reactions at a final concentration of 100 nM of the fluorophore-bearing strand.

### Endpoint visual readout and smartphone imaging of LAMP-OSD assays

LAMP-OSD assays intended for visual readout and smartphone imaging were assembled in 0.2 ml optically clear thin-walled tubes with low auto-fluorescence (Axygen, Union City, CA, USA). Following 90 min of amplification at 65 °C, LAMP-OSD reactions were incubated at 95 °C for 1 min followed by immediate transfer to room temperature and fluorescence imaging. Images were acquired using an unmodified iPhone 6 and an UltraSlim-LED transilluminator (Syngene, Frederick, MD, USA). In some experiments, our previously described in-house 3D-printed imaging device [42] was used for LAMP-OSD fluorescence visualization and smartphone imaging. Briefly, this device uses Super Bright Blue 5 mm light emitting diodes (LED) (Adafruit, New York, NY, USA) to excite OSD fluorescence. Two cut-to-fit layers of inexpensive >500 nm bandpass orange lighting gel sheets (Lee Filters, Burbank, CA, USA) on the observation window filter the OSD fluorescence for observation and imaging.

### Rearing and collection of mosquitoes

*Ae. aegypti, Ae. Albopictus, Culex tarsalis, Cu. quinquefasciatus* (Houston), and *Cu. quinquefasciatus* (Salvador) mosquitoes were reared under conventional conditions in the insectary at the University of Texas medical Branch, Galveston, TX, USA. Four to seven day old mosquitoes were collected and immediately frozen for shipment, storage and subsequent testing. To obtain blood fed insects, *Aedes* mosquitoes were starved for a period of 24 hours then offered a sheep blood meal (Colorado Serum Company, Denver, CO, USA) using a hemotek membrane system (Hemotek). Unfed mosquitoes were separated and mosquitoes that were engorged were collected 24 hours post feeding and processed in the same manner as unfed mosquitoes. For field collections, female mosquitoes were trapped using Fay prince trap (John W. Hock) baited with CO_2_ in Galveston, Texas. 90 mosquitoes were morphologically identified and sorted into three blinded groups that were stored at −20 °C, 4 °C and 37 °C, respectively for up to 3 weeks prior to LAMP-OSD analysis.

### LAMP-OSD analysis of mosquitoes

For LAMP-OSD analysis individual mosquitoes were prepared either in 1 cc syringes or in 1.5 ml microcentrifuge tubes as follows. In-syringe preparation: The plunger was removed from a 1 cc syringe and a 0.5 µM pore size 1/8^th^ inch diameter frit (catalog # 59037, Sigma-Aldrich, St. Louis, MO, USA) was placed inside the syringe. A single mosquito was placed on top of the frit and macerated thoroughly using the syringe plunger. 100 µl of water was aspirated into the syringe to fully re-suspend the macerated mosquito prior to evicting this mosquito-containing water from the syringe into a microcentrifuge collection tube. A 2 µl aliquot of this sample was directly tested by LAMP-OSD assays. In-tube preparation: A single mosquito was placed in a 1.5 ml microcentrifuge tube and manually macerated using a disposable micropestle (Fisherbrand™ RNase-Free Disposable Pellet Pestles, Cat # 12-141-364, Fisher Scientific, Hampton, NH, USA). Each macerated mosquito was resuspended in 100 µl water. A 2 µl 1:10 diluted aliquot of this mosquito sample was directly assessed by LAMP-OSD analysis.

### Statistical analysis

The paired results of morphological identification and LAMP-OSD analysis were compared using 2×2 contingency tables. Sensitivity or true positive rate was calculated by using the formula TP/(TP+FN) where TP are true positive samples, and FN are false negative samples. Specificity or true negative rate was calculated using the formula TN/(TN+FP) where TN are true negative samples and FP are false positive samples.

## Results

### Development of visually read LAMP-OSD assays to identify mosquito species and *Wolbachia* infection

LAMP uses two inner (FIP and BIP) and two outer (F3 and B3) primers specific to six consecutive target sequences (B3, B2, B1, F1c, F2c and F3c) (**S2 Fig**) [47]. *Bst* DNA polymerase extends these primers by strand displacement DNA synthesis to form 10^9^ to 10^10^ copies of concatemerized amplicons with loops between the F1 and F2, B1 and B2, F1c and F2c, and B1c and B2c regions. We use an additional fifth primer that binds to one of these loop regions and accelerates amplification [41]. OSD probes, with blocked 3’-ends that prevents spurious signaling from polymerase-mediated extension, hybridize to the second loop region (**S2 Fig**).

To enable molecular identification of *Ae. aegypti* mosquitoes, we designed a smartphone-imaged LAMP-OSD assay to amplify and detect a signature sequence in the mitochondrial cytochrome c oxidase I (*coi*) gene. Each cell has multiple mitochondria and hence several hundred copies of the *coi* gene, which should enable detection from a very small amount of sample. Moreover, mitochondrial *coi* gene sequences are commonly used as barcodes for molecular identification of animal species including distinction of mosquito species; our chosen *coi* signature sequence was assigned to *Ae. aegypti* when queried against the Barcode of Life Data Systems (BOLD; http://www.boldsystems.org/index.php) *coi* signature sequence database [48–51].

We developed a second visually read LAMP-OSD assay targeting the *Wolbachia* surface protein (*wsp*) gene to identify *Wolbachia*-infected insects. The *wsp* gene is widely used as a marker for strain typing and screening for infected insect vectors [28, 52]. We engineered our *wsp* LAMP outer and inner primers to be complementary to, and hence amplify, two *Wolbachia* strains, the *w*AlbB and the closely related *w*Pip (**S3 Fig**). We deliberately designed our assay to detect both strains in order to ensure that we could assess field-collected mosquitoes irrespective of temporal and spatial variation in relative abundance of *w*AlbB-infected *Ae. albopictus* and *w*Pip-infected *Cu. quinquefasciatus* mosquitoes in our collection area [53, 54]. Significant nucleic acid sequence variation should prevent amplification of the *wsp* gene from all other *Wolbachia* groups (**S3 Fig**). To enable transduction of both *w*AlbB and *w*Pip *wsp* LAMP amplicons to visible fluorescence we designed an OSD probe that is specific to an identical loop sequence present in both amplicons.

With a single endpoint visual ‘yes/no’ readout of OSD fluorescence (either directly observed or imaged using cellphone camera), the *Ae. aegypti coi* LAMP-OSD assay could reliably identify the presence of as few as 4 copies of synthetic target DNA (**Fig 1**). Similarly, the cellphone-imaged *wsp* LAMP-OSD assay produced bright visible fluorescence when presented with only 40 copies of its target sequence. In the absence of target DNA, neither assay generated spurious signal.

**Fig 1.**
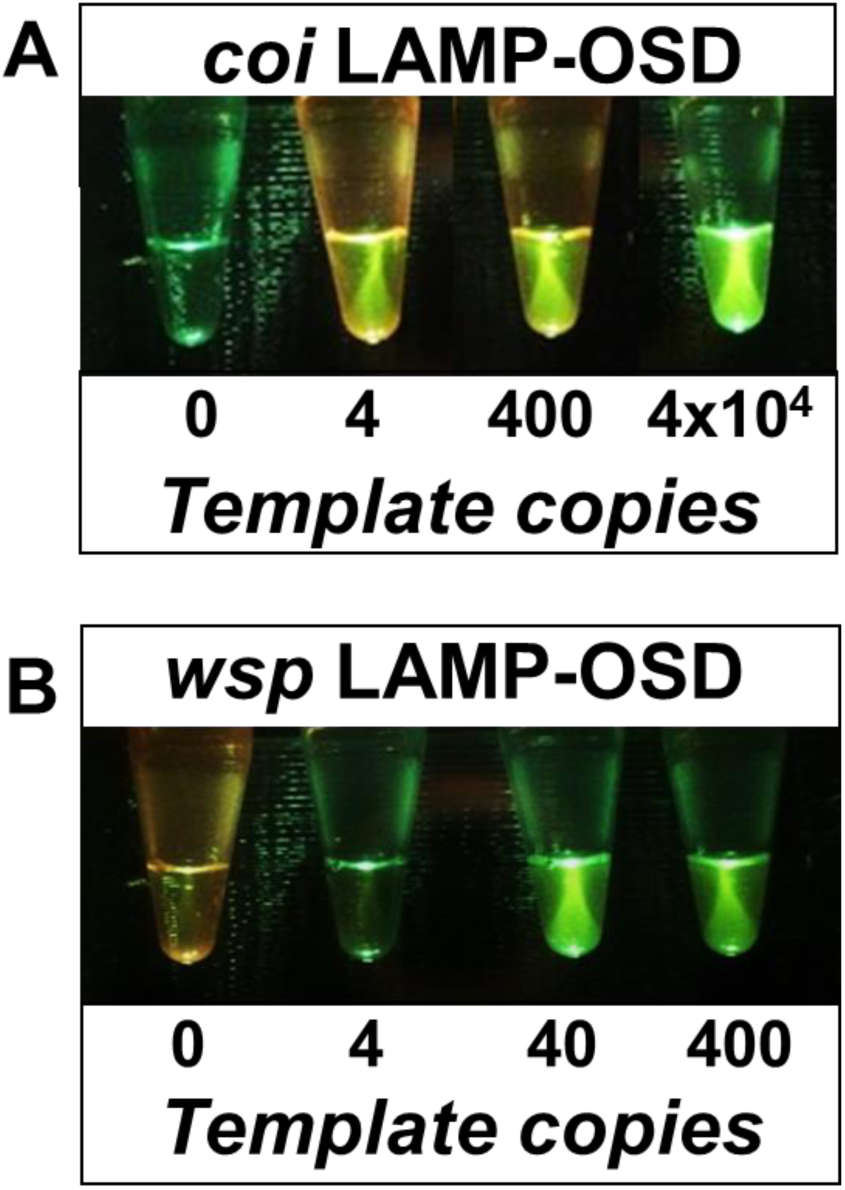
Detection of *Ae. aegypti coi* gene and *Wolbachia w*AlbB *wsp* gene using visually read LAMP-OSD assays. Indicated copies of recombinant plasmids bearing *coi* or *wsp* target sequences were amplified by the *coi*-specific **(A)** or *wsp*-specific **(B)** LAMP-OSD assays, respectively. OSD fluorescence was imaged at endpoint using a cellphone.

### Direct LAMP-OSD analysis of crudely crushed mosquitoes

Our next goal was to demonstrate the ability of these LAMP-OSD assays to detect naturally occurring target sequences in mosquitoes. At the same time, we wanted to ensure that minimally processed samples would be compatible with our detection platform in order to facilitate rapid in-field vector testing with fewest instruments and user-required steps.

Therefore, as an initial approach, we developed the ‘in-syringe’ method for rapid sample preparation wherein individual mosquitoes were crushed inside 1 cc syringes using the syringe plunger as a pestle. A small chromatography column frit placed inside the syringe served as a pedestal that aided maceration and removed larger particulates when the macerated sample was re-suspended in water and recovered. Small portions (up to 8% of a LAMP-OSD reaction) of these macerated samples were added directly to LAMP-OSD reactions, which were then incubated for 90 min at 65 °C to initiate and sustain amplification.

Endpoint visual examination of these assays for the presence or absence of OSD fluorescence revealed that our visually read LAMP-OSD system is compatible with direct analysis of crudely processed mosquitoes (**Fig 2**). The *coi* LAMP-OSD assay generated bright fluorescence readily distinguishable from sample auto-fluorescence when seeded with crudely prepared *Ae. aegypti* mosquitoes. In contrast, closely related *Ae. albopictus* failed to instigate false positive signal. Similarly, the *wsp* LAMP-OSD assay generated bright fluorescence in response to *Ae. albopictus* and *Cu. quinquefasciatus* mosquitoes, which are naturally infected with *w*AlbB and *w*Pip *Wolbachia*, respectively, but remained negative in the presence of unrelated *Wolbachia w*Mel and uninfected mosquitoes (**Fig 2 and S4 Fig**).

**Fig 2.**
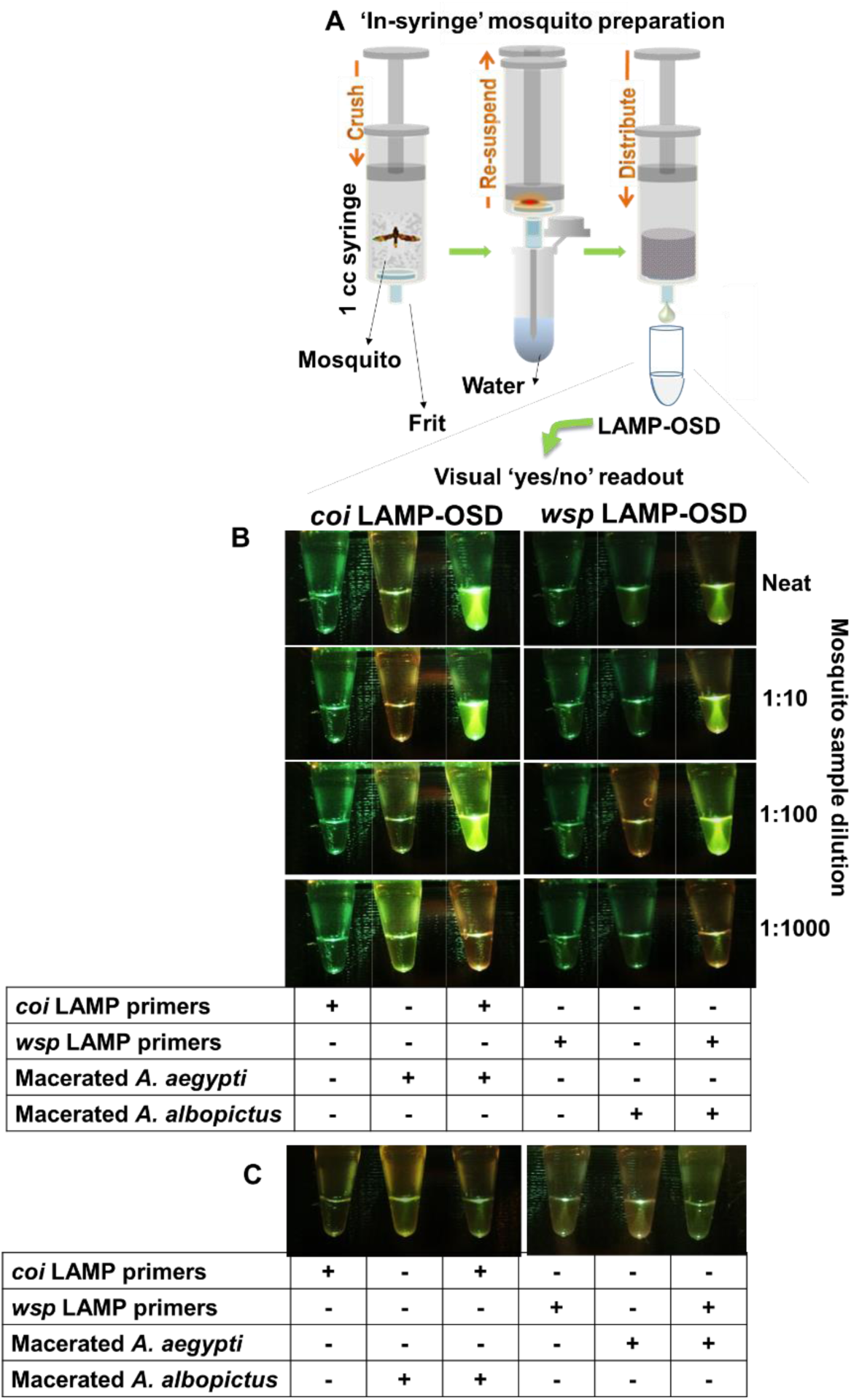
Visually read LAMP-OSD analysis of crudely processed mosquito samples. **(A)** Schematic depicting rapid preparation of crude mosquito samples for direct LAMP-OSD analysis by performing in-syringe individual mosquito maceration. **(B)** Analysis of individual *Ae. aegypti* and *w*AlbB *Wolbachia*-infected *Ae. albopictus* mosquitoes by *coi* and *wsp* LAMP-OSD assays, respectively. 2 µL of mosquito samples were either added directly to LAMP-OSD assays or subjected to 10-fold serial dilution in water prior to introduction in LAMP-OSD reactions. LAMP-OSD reactions without any macerated mosquito samples served as negative controls. Assays lacking LAMP primers but containing macerated mosquito samples served as controls for monitoring background fluorescence. **(C)** Specificity analysis of *coi* and *wsp* LAMP-OSD assays. The *coi* LAMP-OSD assay was challenged with 2 µL of *Ae. albopictus* in-syringe prepared crude mosquito sample. Similarly, the *wsp* LAMP-OSD assay was challenged with *Ae. aegypti* sample. In all experiments, OSD fluorescence was imaged at endpoint using a cellphone.

These results indicate that the *Wolbachia wsp* and *Ae. aegypti coi* LAMP-OSD assays are able to specifically amplify and signal the presence of their target DNA directly from crudely crushed mosquito samples without requiring any extraction and purification of nucleic acids. Furthermore, the large burden of non-specific nucleic acids as well as other molecular and macroscopic components present in a crude mosquito sample did not compromise signal accuracy. We also confirmed the absence of significant inhibition of amplification and signaling by recapitulating the detection limit of synthetic DNA targets in a background of crude non-specific mosquito sample. The *coi* and *wsp* LAMP-OSD assays could detect 4 and 40 target copies, respectively, even in the presence of 8% reaction volume of crude mosquito sample (**S5 Fig)**.

### Testing blood-fed mosquitoes with LAMP-OSD

Mosquitoes feeding on blood meals have been reported to engorge on 1 nL to as much as 6 µL of blood [55]. It is conceivable that the blood meal might confound visual LAMP-OSD fluorescence analysis by contributing auto-fluorescence. To ascertain compatibility of visually read LAMP-OSD with direct analysis of crudely prepared blood-engorged mosquitoes, we challenged both *coi* and *wsp* LAMP-OSD assays with crude in-syringe preparations of blood engorged *Ae. aegypti* and *Ae. albopictus* mosquitoes. Mosquitoes that had recently consumed a blood meal could be directly analyzed by visual LAMP-OSD without diminution of signal to noise ratio (**Fig 3**).

**Fig 3.**
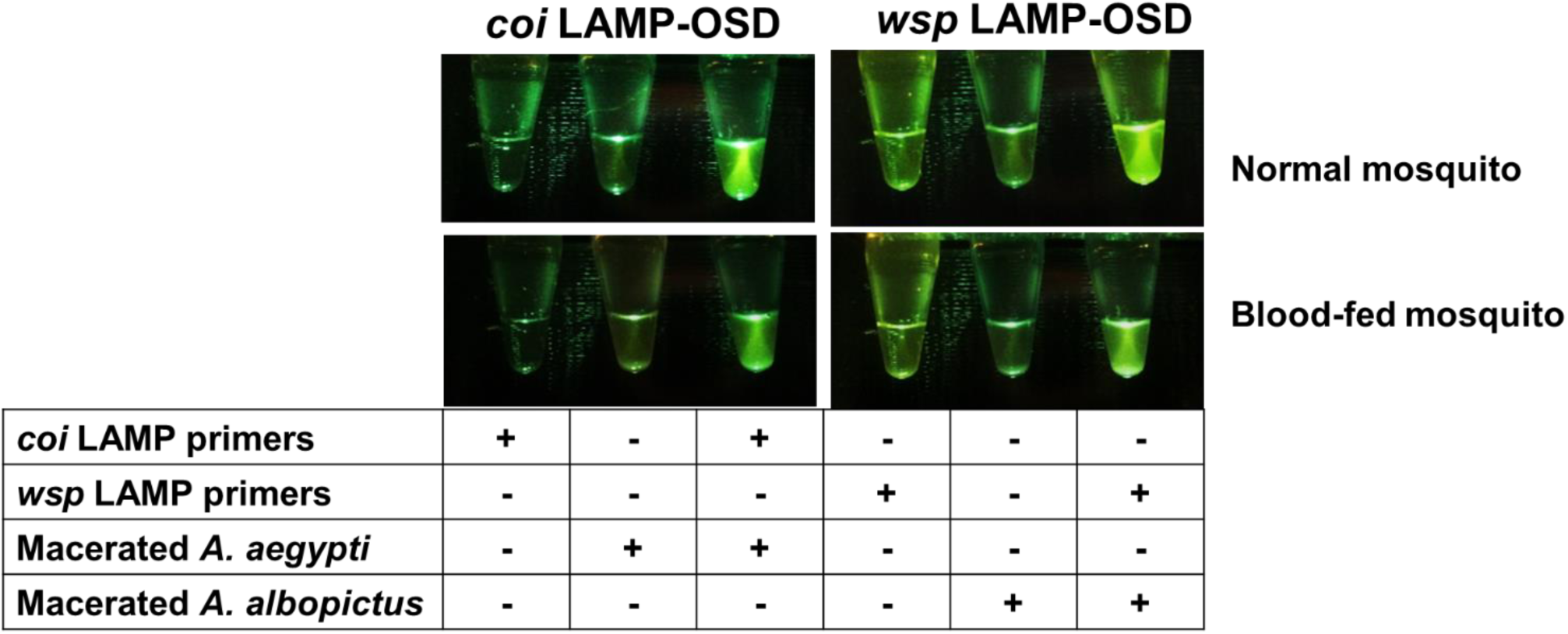
Effect of blood meal on LAMP-OSD analysis of crudely prepared individual mosquitoes. Individual normal or blood-fed *Ae. aegypti* and *Ae. albopictus* mosquitoes were prepared by the in-syringe method. 2 µL of these mosquito samples were analyzed by *coi* and *wsp* LAMP-OSD assays. LAMP-OSD reactions without any macerated mosquito samples served as negative controls. Assays lacking LAMP primers but containing macerated mosquito samples served as controls for monitoring background fluorescence. LAMP-OSD fluorescence was imaged at endpoint using a cellphone.

### Analysis of blinded field-caught mosquitoes with cellphone-imaged LAMP-OSD assays

To validate assay performance under more rigorous conditions, we challenged the LAMP-OSD system with a blinded set of 90 field-caught mosquitoes comprised of *Ae. aegypti*, *Ae. albopictus*, and *Ochlerotatus* species. The mosquitoes were divided into three groups of 30 individuals that were stored without desiccant at −20 °C, 4 °C, or 37 °C for 1, 2, or 3 weeks prior to testing. To reduce mosquito processing cost, footprint, and time for this large study, we further simplified sample preparation requirements by optimizing the “in-tube” mosquito preparation method wherein each mosquito was crushed with a micropestle directly in a microcentrifuge tube followed by resuspension in water and introduction in a LAMP-OSD reaction.

The visually read *coi* LAMP-OSD assay demonstrated an overall sensitivity (true positive rate) of 97% and specificity (true negative rate) of 98% when compared to morphological typing of field-caught mosquito species (**Fig 4**, **S6, S7, S8**). On closer inspection of the data, it is evident that even after three weeks of mosquito collection and storage at temperatures as high as 37 °C the *coi* LAMP-OSD assay was correctly able to identify 29 out of 30 *Ae. aegypti* mosquitoes. The single mosquito that the LAMP-OSD assay failed to identify had been stored at 37 °C for a week prior to testing. We ruled out lack of amplifiable nucleic acids or their incompatibility with *coi* LAMP primers and OSD probe by PCR amplifying the relevant *coi* LAMP target and verifying its sequence.

**Fig 4.**
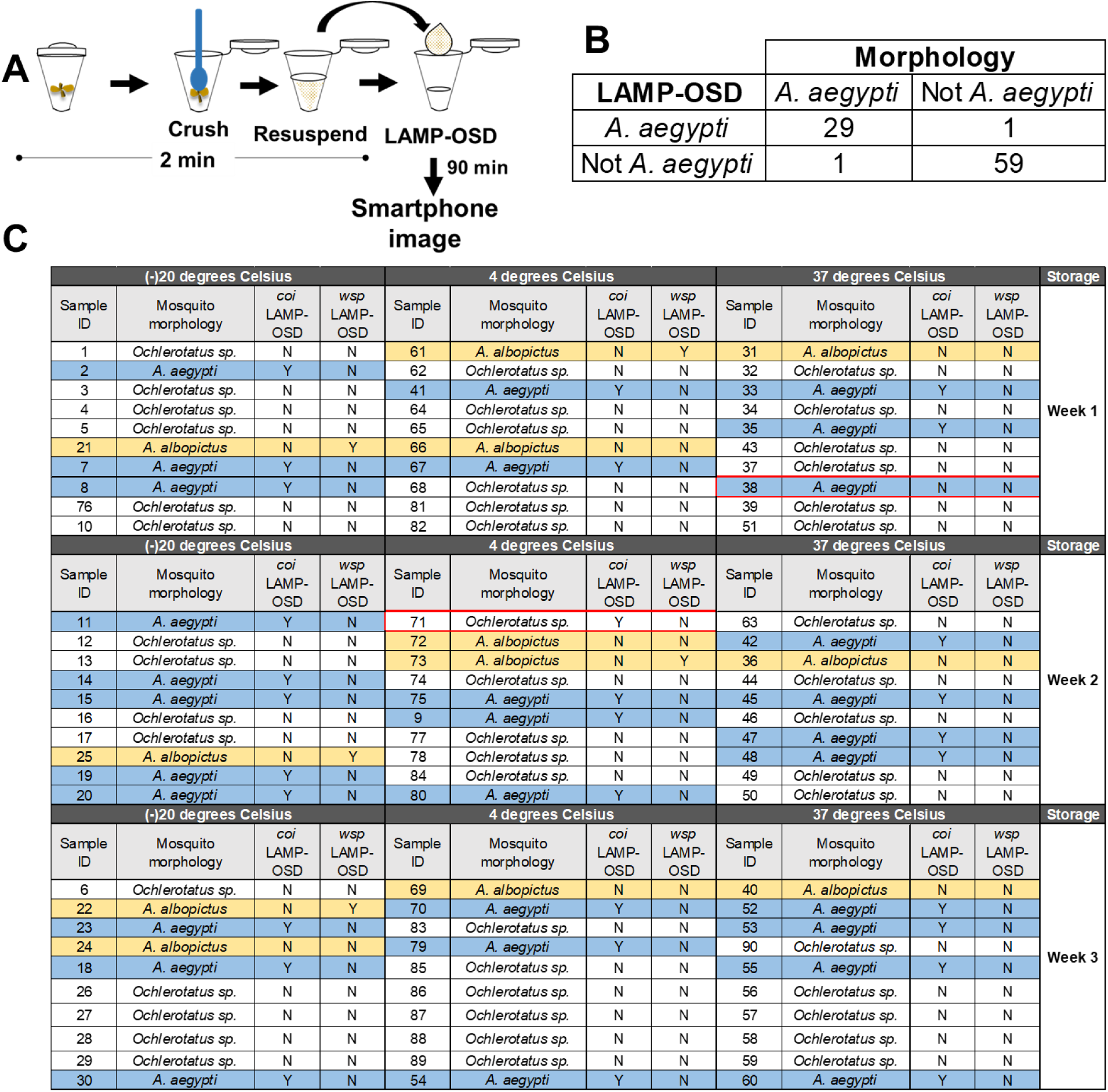
Blinded LAMP-OSD analysis of field-caught mosquitoes. Mosquitoes stored for up to three weeks at the indicated temperatures were prepared by ‘in-tube’ crude processing. 2 µL of a 1:10 dilution of each mosquito sample was analyzed by *wsp* and *coi* LAMP-OSD assays. OSD fluorescence was imaged at endpoint using a cellphone. **(A)** Schematic depicting ‘in-tube’ method for rapid preparation of field-caught mosquitoes for LAMP-OSD analysis. **(B)** Comparison of *coi* LAMP-OSD results with morphological identification using 2×2 contingency table. **(C)** Tabulation of *coi* and *wsp* LAMP-OSD readout for each mosquito in the study. *Ae. aegypti, Ae. albopictus* and *Ochlerotatus* species are highlighted in blue, orange and white, respectively. ‘Y’ indicates a positive LAMP-OSD signal (bright fluorescence). ‘N’ indicates absence of any fluorescence signal in the LAMP-OSD reaction. Red boxes highlight LAMP-OSD tests that generated false positive or false negative outcomes.

Of the 60 non-*Ae. aegypti* mosquitoes analyzed by *coi* LAMP-OSD, only one mosquito generated a false positive signal. Sequence analysis of its *coi* gene ruled out mis-firing of the *coi* LAMP-OSD assay. It is possible that this LAMP assay was inadvertently contaminated with a small amount of a pre-formed *Ae. aegypti* amplicon.

The *Wolbachia w*AlbB/*w*Pip *wsp* LAMP-OSD assay did not generate a positive signal from any non-*Ae. albopictus* mosquito. This is expected since natural populations of *Ae. aegypti* and most *Ochlerotatus* species are not infected with *Wolbachia* [56, 57]. However, ability of the *wsp* assay to identify *Wolbachia* infection was influenced by the storage temperature of mosquitoes. The *wsp* LAMP assay could readily identify *Wolbachia* infection in 3 out of 4 *Ae. albopictus* mosquitoes stored at −20 °C for as long as 3 weeks. PCR analysis of the *wsp*-negative mosquito using previously described primers (81F and 691R) and protocols [28] did not produce amplicons suggesting that this individual was likely uninfected or had *Wolbachia* levels below the levels detectable by PCR. As the storage temperature was increased the frequency of *Wolbachia* detection dropped. While 40% of *Ae. albopictus* mosquitoes stored at 4 °C gave a positive *wsp* LAMP-OSD signal, none of the *Ae. albopictus* mosquitoes kept at 37 °C for even as little as 1 week were *wsp* positive. All mosquitoes that failed to generate a signal by LAMP also failed to produce *wsp* PCR amplicons. Since, 95-99% of *Ae. albopictus* mosquitoes in the wild are typically found to be infected with *Wolbachia* [58], these results are suggestive of nucleic acid deterioration in mosquitoes upon storage at high temperature.

## Discussion

Mosquito control strategies that rely on the introduction of *Wolbachia* are now being deployed around the world [7–9], and the surveillance of efficacy and spread require agile, field-based methods for both mosquito and symbiont detection. Unfortunately, currently available tools for mosquito diagnostics have several shortcomings. Morphological identification methods are inherently low throughput, require extensive technical expertise, and cannot also readily identify pathogens or biocontrol agents such as *Wolbachia*. Spectroscopy, such as near infrared spectroscopy [59] and Fourier transform infrared spectroscopy [60], allow identification of mosquito species, *Wolbachia,* and pathogens, but require expensive instruments and expertise that are generally incompatible with low-resource settings. Immunoassays can detect pathogen-carrying vectors but are insensitive and cannot also identify vector species. Nucleic acid amplification methods could potentially look at both vector and sybmiont sequences, but are heavily reliant on PCR with the ensuing encumbrances of expensive instruments and trained operators, again precluding widespread use.

Isothermal nucleic acid amplification assays would facilitate field-based vector monitoring, but most reported approaches rely on nucleic acid purification and non-specific readout, and thus suffer from laborious setup and the risk of false positives [61–63]. Probe-read isothermal methods such as the recombinase polymerase assay (RPA) are more reliable but still require expensive and proprietary reaction formulations and probes, which limits their flexibility and versatility in assay engineering. Furthermore, most RPA applications for vector diagnostics [64] also depend on extensive sample processing and nucleic acid purification prior to amplification.

These drawbacks led us to develop a simpler, more robust field-deployable assay based on loop-mediated isothermal amplification (LAMP) that can identify both mosquito species and specific *Wolbachia* strains in mosquitoes. While LAMP assays have previously been developed for *Wolbachia* detection by targeting the 16S rRNA gene for amplification [61], these assays required extraction and purification of DNA prior to assay, and used non-specific readouts that were highly prone to false positive signals.

To overcome these barriers to the use of LAMP, we have previously adopted methods that originated in the field of nucleic acid computation: the use of strand exchange reactions that initiate at complementary single-stranded ‘toeholds’ and progresses via branch migration. The base-pairing predictability and programmability of strand exchange kinetics promotes the construction of exquisitely sequence-specific oligonucleotide strand displacement (OSD) probes for LAMP amplicons (**S2 Fig**), thereby greatly reducing the detection of non-specific amplification background [39, 41, 42]. For instance, we recorded positive *coi* LAMP-OSD signals from field-caught *Ae. aegypti* mosquitos but did not detect signal from closely related *Ae. albopictus* and *Ochlerotatus* species mosquitoes. Similarly, the *wsp* assay detected *w*AlbB and *w*Pip, as expected, but not *w*Mel *Wolbachia*.

Strand exchange circuits have the additional advantage that they can be used to embed algorithms and act as ‘matter computers’ [35–37, 65]. For example, strand exchange transducers can logically integrate multiple analytes; transform nucleic acids to glucose and human chorionic gonadotrophin (hCG); adapt readout to beads, paperfluidics, glucose meters, pregnancy test strips, and cellphones, and allow target copy number estimation using a single endpoint yes/no readout of presence or absence of signal above an adjustable threshold [66–69]. In the current instance, we deliberately designed our *wsp* LAMP-OSD assay to ‘compute’ the presence of both *w*AlbB and *w*Pip in order to increase our odds of finding infected field-caught mosquitoes. However, the dependence of strand exchange efficiency on toehold binding strength [39] can be exploited to engineer yes/no distinctions between strain-specific single nucleotide polymorphisms [39], and the same dual *wsp* assay could be rendered strain-specific by simply substituting an OSD reporter specific to an alternate polymorphic loop sequence (**S3 Fig**). This might be advantageous for strain discrimination if double infections were released during vector control measures [70, 71].

By using our one-pot LAMP-OSD assay, macerated mosquito homogenates could be directly analyzed and ‘yes/no’ visual readouts could be quickly ascertained with a cell phone in the field without the requirement for laboratory equipment or technically training. Moreover, since our assays can accurately analyze mosquitoes several days after capture – the *coi* LAMP-OSD assay could for example identify mosquitoes after 3 weeks at 37 °C without desiccant – mosquitoes from remote collection outposts can potentially be analyzed even after delayed retrieval. The flexibility of assay timing is further accommodated by the fact that lyophilized LAMP-OSD reaction mixes that can be stored and deployed without cold chain [42]. These combined features make our assay platform the best tool to date for expanding vector surveillance to resource poor settings [72], especially in that the ease of use should allow minimally trained citizen scientists to participate in otherwise sophisticated public health monitoring operations in the field.

The development efforts that we have put into LAMP-OSD should now allow it to be generalized to screening other microbes or mosquito phenotypes in field settings. For example, LAMP-based assays have been developed to identify pathogens transmitted by mosquitoes and insecticide resistance alleles [73–76], but these rely on purified nucleic acid as templates and non-specific readout whereas our method functions with mosquito homogenate and ensures accuracy using unique sequence-specific strand exchange probes. In addition, this technology could be used to identify gut microbes and insect-specific viruses associated with mosquitoes, which is of growing interest given that it is becoming clear that the microbiome can shape vector competence for human pathogens [77–79]. Overall, we have demonstrated a versatile nucleic acid diagnostic platform for rapid and accurate analyses of both insect vectors and symbionts, and that can now be further configured for additional applications.

## Acknowledgements

We would like to thank Dr. Holly Wichman (Department of Biological Sciences, Center for Modeling Complex Interactions, University of Idaho, Moscow, ID 83844-3051) for the generous gift of *w*Mel *Wolbachia*-infected *Drosophila* fruit flies. We would also like to thank the UTMB insectary core for providing the lab-reared mosquitoes.

## Supporting information

**S1 Table. Primers, probes, and target sequences for A. aegypti coi and Wolbachia wAlbB/wPip wsp LAMP-OSD assays.**

**S2 Fig. LAMP-OSD schematic.** LAMP uses 2 inner (FIP and BIP) and 2 outer (F3 and B3) primers specific to 6 blocks of target sequences designated as B3, B2, B1, F1c, F2c and F3c. F2 sequence in FIP (F1c-F2) initiates amplification by Bst DNA polymerase (Stage I). F1c sequence in FIP self-primes subsequent amplification. Similarly, BIP (B1c-B2) initiates DNA synthesis by binding to B2c. F3 and B3 primer-initiated DNA synthesis displaces preceding inner primer-initiated strands, which serve as templates for primer-initiated strand displacement DNA synthesis (Stage II). 3′-ends of the resulting single-stranded, dumbbell-shaped amplicons (Stage III) are extended by Bst polymerase to form hairpins (Stage IV). Inner primers hybridize to the single-stranded loops and initiate another round of strand displacement synthesis that opens the original hairpin to form a concatemerized amplicon containing a self-priming 3′-end hairpin (Stage V). The ensuing continuous amplification (initiated both by new inner primers and by self-priming) generates increasingly complex, double-stranded concatameric amplicons containing self-priming hairpins and single-stranded loops to which the OSD probe hybridizes. “c”: denotes complementary target sequences. F and Q on the OSD denote fluorophore and quencher, respectively. OSD probe is denoted in terms of numbered domains, each of which represents a short fragment (usually <12 nt) of DNA sequence in an otherwise continuous oligonucleotide strand. Single stranded toeholds are numbered in red. Complementarity between numbered OSD domains is denoted by a single prime symbol.

**S3 Fig. Alignment of *wsp* sequences from different *Wolbachia* strains.** “Current *wsp* OSD” refers to the *wsp* OSD probe used in the present study that binds to the loop sequence between the F1 and F2 target regions. It would not distinguish the closely related *w*AlbB and *w*Pip strains. The “*w*AlbB vs *w*Pip OSD”, which would bind to the loop region between the B1 and B2 regions of our current *wsp* LAMP assay would allow discrimination of *w*AlbB and *w*Pip strains due to specificity of interaction with the three highlighted polymorphic positions. The *wsp* sequences of the remaining *Wolbachia* strains are significantly different from *w*AlbB *wsp* sequence. Hence, the *wsp* LAMP-OSD assay described here will not detect these strains.

**S4 Fig. Detection of *Wolbachia w*Pip using *wsp* LAMP-OSD assay.** *Wolbachia w*Mel-infected *Drosophila melanogaster* **(A)**, *Wolbachia* uninfected *Culex tarsalis* **(B)**, *Wolbachia w*Pip infected *Culex quinquefasciatus* (Houston) **(C)** and *Culex quinquefasciatus* (Salvador) **(D)** were analyzed using the *wsp* LAMP-OSD assay. 2 µL of crudely crushed individual insect samples (crushed individual fruit flies were resuspended in 20 µL water) were subjected to LAMP amplification for 90 min. OSD fluorescence was imaged at endpoint using a cellphone.

**S5 Fig. Effect of crude mosquito sample on the detection limit of visually-read LAMP-OSD.** Indicated copies of recombinant plasmids bearing *coi* or *wsp* target sequences were amplified by the *coi*-specific **(A)** or *wsp*-specific **(B)** LAMP-OSD assays, respectively. 8% volume of all LAMP-OSD reactions was composed of crudely ‘in-syringe’ prepared non-specific mosquito sample. OSD fluorescence was imaged at endpoint using a cellphone.

**S6 Fig. Blinded LAMP-OSD analysis of field-caught mosquitoes.** Mosquitoes stored for one week at the indicated temperatures were prepared by ‘in-tube’ crude processing. 2 µL of a 1:10 dilution of each mosquito sample was analyzed by *coi* LAMP-OSD (A) and by *wsp* LAMP-OSD (B) assays. OSD fluorescence was imaged at endpoint using a smartphone.

**S7 Fig. Blinded LAMP-OSD analysis of field-caught mosquitoes.** Mosquitoes stored for two weeks at the indicated temperatures were prepared by ‘in-tube’ crude processing. 2 µL of a 1:10 dilution of each mosquito sample was analyzed by *coi* LAMP-OSD (A) and by *wsp* LAMP-OSD (B) assays. OSD fluorescence was imaged at endpoint using a smartphone.

**S8 Fig. Blinded LAMP-OSD analysis of field-caught mosquitoes.** Mosquitoes stored for three weeks at the indicated temperatures were prepared by ‘in-tube’ crude processing. 2 µL of a 1:10 dilution of each mosquito sample was analyzed by *coi* LAMP-OSD (A) and by *wsp* LAMP-OSD (B) assays. OSD fluorescence was imaged at endpoint using a smartphone.

